# Mass Spectrometric Profiling of Microbial Polysaccharides Using Laser Desorption/Ionization – Time-of-Flight (LDI-TOF) and Liquid Chromatography/Mass Spectrometry (LC/MS): A Novel Method for Structural Fingerprinting and Derivatization

**DOI:** 10.1101/2025.05.13.653920

**Authors:** Lucia Dadovska, Veronika Paskova, Petr Novak, Jaroslav Hrabak

## Abstract

Over the last two decades, matrix-assisted laser desorption/ionization time-of-flight mass spectrometry (MALDI-TOF MS) has been introduced into the routine diagnostic practice of microbiological laboratories for the rapid taxonomic identification of bacteria and yeasts. However, a method that effectively identifies microbes directly from clinical samples using MALDI-TOF MS has not yet been found. One of the promising targets is microbial polysaccharides, which are abundant structures in bacterial and fungal cells. Their rapid and inexpensive analysis, however, is complicated. This study focused on detecting microbial polysaccharides, such as lipopolysaccharides, using MALDI-TOF MS and liquid chromatography-tandem mass spectrometry (LC-MS). We developed a method for fingerprinting polysaccharides by acid hydrolysis and enzymatic digestion. The mono- and oligosaccharides are then derivatized with a newly developed probe (vanillyl pararosaniline, HD ligand), enabling efficient ionization without the use of the MALDI matrix. The esterification of hydroxyl groups by formic acid was also optimized for precise analysis of the saccharides. The method was validated using several saccharides as well as *Escherichia coli* lipopolysaccharides (O26:B6, O55:B5, and O111:B4). Derivatization using HD ligand also allows the detection of structures containing amines and phosphate groups in positive ion mode. We also optimize the method using crude bacteria (*Escherichia coli*, *Salmonella enterica*, *Shigella dysenteriae*, *Shigella boydii*, *Shigella flexneri*, and *Legionella pneumophila*). This approach opens the possibility of directly detecting microbial polysaccharides from clinical specimens. LDI-TOF MS (without a matrix) also allows specific detection of molecules of interest with suppression of the background signal.

**Scheme 1.**
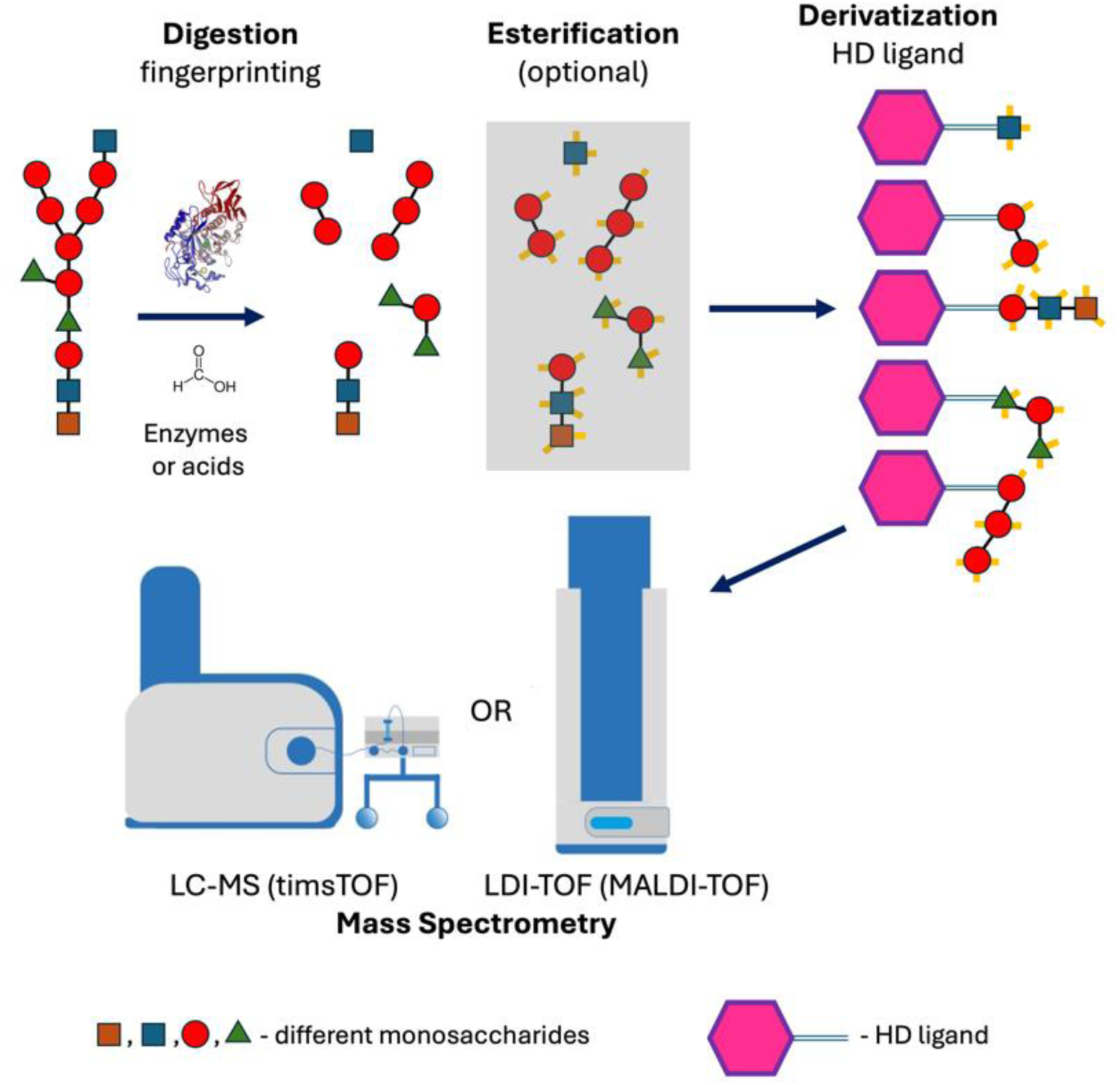
Analysis of polysaccharides by mass spectrometry after digestion and derivatization using a self-ionizable ligand (HD ligand).

## INTRODUCTION

Many innovative applications of matrix-assisted laser desorption/ionization time-of-flight mass spectrometry (MALDI-TOF MS) and liquid chromatography coupled with mass spectrometry (LC-MS) have emerged in clinical diagnostics to detect biomarkers and biomolecules, advancing personalized medicine [1, 2].

In diagnostic microbiology, MALDI-TOF MS became a cornerstone in the rapid taxonomic identification of bacteria and fungi [2, 3]. Similarly, applications for antibiotic resistance determination have also been developed and validated. Among them, beta-lactamase activity determination detecting the molecular mass changes of indicator beta-lactam antibiotics (e.g., carbapenems) has been developed and validated for use in routine laboratory practice [4]. Another method allows the detection of polymyxin resistance using analysis of lipid A of lipopolysaccharides and their structural changes (e.g., the addition of phosphoethanolamine or 4-amino-L-arabinose) [5].

As MALDI-TOF MS provides efficient and rapid species identification, there is a key issue whether the method can be used for epidemiological typing directly from spectra obtained directly from a whole-cell lysate. So far, no general typing algorithm has been proposed, but only specific peaks representing significant epidemiological markers have been identified in some species [3]. Artificial intelligence for spectra analysis has recently been described as a promising tool for predicting antibiotic resistance and epidemiological typing [6].

Despite the use of artificial intelligence methods for analyzing large-scale data, MALDI-TOF MS should be considered a biochemical tool allowing precise analysis of molecules based on their molecular weight and fragmentation characteristics. Thus, we believe that the scientific community should not resign itself to the exact identification of detected molecules/peaks. For such an analysis, it is usually insufficient to analyze crude bacterial extract without further processing, i.e., specific extraction and enhancement of MALDI-TOF MS-based ionization.

Despite the commonly used taxonomic applications in routine laboratories, methods for identifying microbes directly from clinical specimens have not yet been successfully developed [3]. Such applications, however, are limited by a poor sensitivity of MALDI-TOF MS and by masking the proteins of interest by other abundant molecules with a high ionization ability. These challenges can be addressed by (a) concentrating microbes or microbial proteins in the sample and removing host proteins, (b) detecting other molecules (e.g., lipids or polysaccharides) that can be explicitly extracted from the sample, and (c) using LC-MS.

Identification of microbial polysaccharides represents a challenge in clinical microbiology as those structures can be used for (a) rapid and cheap epidemiological typing (e.g., *Salmonella enterica*, *Shigella* spp., *Escherichia coli*), (b) development of polysaccharide vaccines and testing their efficiency, and (c) direct identification of microbes from clinical specimens. The latter option is challenging as microbial saccharides represent relatively stable molecules that can be directly detected in several clinical specimens (e.g., blood, urine), allowing rapid identification of a causative agent of the infection and thus increasing therapeutic efficiency [7, 8, 9].

As microbial polysaccharides usually contain >10 basic units, their molecular weight is highly above the efficient ability of MALDI-TOF MS or LC-MS to detect them directly. Therefore, the native molecules must be specifically cleaved into smaller units, providing a specific molecule fingerprint. For that purpose, the glycosidic bond is the common target. Hydrolysis of that bond, however, is challenging due to its stability in some biological structures. Several procedures of polysaccharide fingerprinting, including chemical cleavage by strong acids or Fenton’s reaction and enzymatic disruption, have been developed and optimized [10, 11, 12].

Even if monosaccharides or oligosaccharides are available, their ionization using conventional mass spectrometry techniques is complex. Compared to peptides and proteins, those molecules are generally more complicated to ionize and transfer to the vapor phase [12]. Therefore, the typical approach is to derivatize mono and oligosaccharides before mass spectrometry. For that purpose, several methods have been proposed, including the label-assisted laser desorption/ionization approach [13].

Here, we present a novel method for enzymatic and acidic hydrolysis of bacterial polysaccharides, their derivatization, and analysis, where the sample is ionized through the addition of a specific ligand, enabling the use of LDI-TOF (Laser Desorption/Ionization Time-of-Flight) MS and LC-MS (Trapped Ion Mobility Spectrometry—TIMS). This approach allows the identification of saccharide fingerprints based on their ion mobility and *m/z*. Additionally, we proposed the esterification of mono- and oligosaccharides by formic acid, which can aid in their identification.

## MATERIALS AND METHODS

### Reagents

For analysis, all chemicals (_D_-glucose, _D_-glucose-1,2-^13^C_2_, glucose-6-phosphate, _L_-glucose, _D_-fructose, _L_-rhamnose, _L_-fucose, _D_-xylose, lactose, maltose, sucrose, raffinose, 5-(hydroxymethyl)furfural), starch, agarose, lipopolysaccharides (*Escherichia coli* O26:B6, *E. coli* O55:B5, *E. coli* O111:B4), enzymes (diastase, α-amylase, β-amylase, lysozyme, chitinase), and other reagents were purchased from Merck (Merck Life Science, Prague, Czech Republic). The greater quantity of vanillyl-rosaniline (HD ligand) was synthesized as 2,3-Dichloro-5,6-dicyano-1,4-benzoquinone salt by Ratiochem (Brno, Czech Republic).

### Instrumentation

Samples for LC-MS were injected and subsequently desalted online using reverse-phase microtrap column (Neo Trap Cartridge 5mm, ThermoFischer Scientific, Prague, Czech Republic) and separated on the reverse-phase analytical column (Bruker TEN, Bruker Daltonics, Bremen, Germany) at 40 °C using Bruker nanoElute 2 HPLC system at flow rate 0.5 µL/min in water/acetonitrile gradient (mobile phase [A] water with 0.1% formic acid; mobile phase [B] acetonitrile with 0.1% formic acid; the gradient started at 10% [B] and reached 50% [B] in 35 min). The eluted analytes were directly analyzed by timsTOF Pro 2 mass spectrometer using a CaptiveSpray for ionization (Bruker Daltonics, Bremen, Germany). Measurement was performed in positive ion mode over the *m/z* range 100 – 1350 in dia-PASEF® mode (0.75 – 1.60 V.s/cm^2^, ramp time 100 ms). Data acquisition and data processing were performed using DataAnalysis 6.1.

LDI-TOF MS was performed using a MALDI Biotyper® Sirius mass spectrometer and rapifleX® MALDI-TOF/TOF system (Bruker Daltonics, Bremen, Germany). The spectra were analyzed using flexAnalysis 4.0 software. For precise identification of molecular mass, the samples were analyzed using a 15T solariX XR FT-ICR mass spectrometer (Bruker Daltonics). Mass spectral data were collected in positive broadband mode over the *m/z* range 150 – 1500, with 1M data points transient and 0.2 s ion accumulation with two averaged scans per spectrum. Data acquisition and data processing were performed using ftmsControl 2.1.0 and DataAnalysis 5.0.

### Synthesis of Vanillyl-Rosaniline Reagent (HD ligand)

Pararosaniline hydrochloride (basic fuchsin) and vanillin were dissolved in methanol to a 100 mmol/L concentration (both reagents). After dissolving, glacial acetic acid was added to a final concentration of 2.5 mol/L and mixed well. Methylpyridine borane dissolved in methanol as a reducing agent was added to the reaction mixture to a final concentration of 10 mmol/L and incubated at 50 °C with shaking for 2 hrs. After incubation, the reaction was diluted by deionized water to a final concentration of methanol of 10 % and directly applied on the NGC Quest Plus System (Bio-Rad, Prague, Czech Republic) equipped with a XSelect® CSH C18 OBDTM preparative column (Waters, Gesellschaft m.b.H., Prague, Czech Republic) at flow rate 0.1 mL/min. After application of the reaction mixture, the column was washed by 50 mL of 5% acetonitrile with 0.1% formic acid. Then water/acetonitrile gradient (mobile phase [A] water with 0.1% formic acid; mobile phase [B] acetonitrile with 0.1% formic acid; the gradient started at 5% [B] and reached 35% [B] in 60 min) was used for sample purification. Two mL fractions were collected and measured using LC-MS after dilution with water 1:10. Fractions showing purified vanillyl-rosaniline (HD ligand) (Figure 1) were dried by a vacuum concentrator.

**Figure 1.**
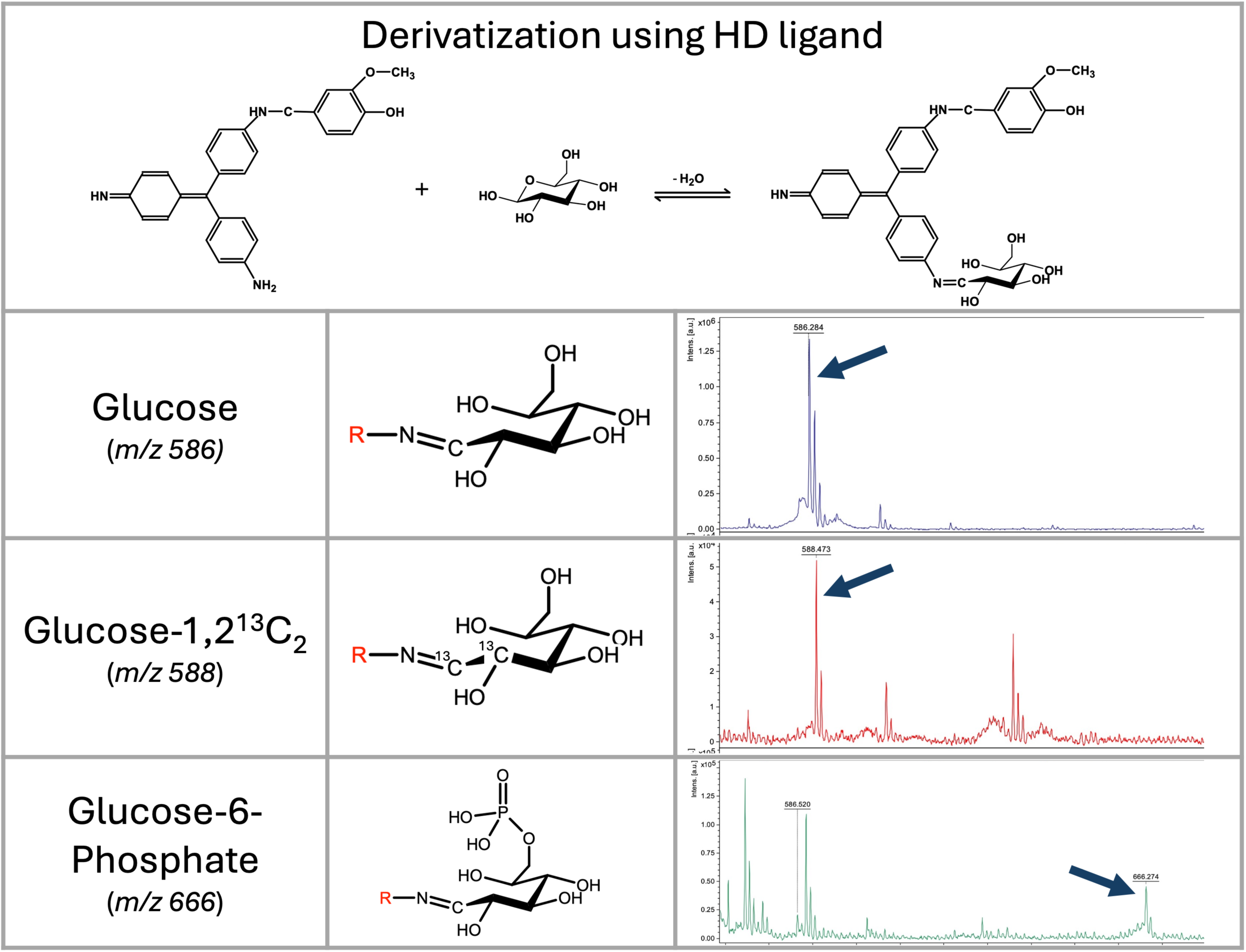
Vanillyl-rosaniline (HD ligand) mechanism of saccharide’s derivatization, and LDI-TOF MS analysis of selected saccharides. Mass spectra of glucose, _D_-glucose-1,2-^13^C_2_, and glucose-6-phosphate derivatized by HD ligand measured on a rapifleX mass spectrometer in linear positive ion mode. Derivatized glucose is visible at *m/z* 586; D-glucose-1,2-^13^C is visible as a signal at *m/z* 588; and glucose-6-phosphate as a signal at *m/z* 666.

### Saccharide’s Derivatization and Analysis

Derivatization was performed in a 1% pyridine buffer, and the pH was adjusted to 4.0 using glacial acetic acid. Saccharides were diluted to a concentration of 5 mmol/L with a reaction buffer. Concentrated HD (250 mmol/L) solution diluted in acetonitrile was added to a final 5 mmol/L concentration. The reaction mixture was incubated at 50 °C for 20 minutes.

For LC-MS, the samples were diluted 1:5 with a pyridine buffer. For LDI-TOF MS, the samples were purified by C18-reversed phase silica gel to avoid the “sweet spot” formation on the MALDI target. Purification was performed in the Eppendorf tube. To 5 mg of C18 reverse phase resin, 10 μL of 30% acetonitrile with 0.1% formic acid was added. After vortexing, 50 μL of the reaction mixture was added, vortexed for 10 s, and centrifuged at 14,000 g. The resin was washed with 1 mL of 5% acetonitrile with 0.1% formic acid. The sample was eluted with 25 μL of 50% acetonitrile with 0.1% formic acid. Two μL of the sample was directly applied to the MALDI target, allowed to dry, and then measured without adding any matrix.

### Enzymatic Digestion of Bacterial Lipopolysaccharides

Two milligrams of bacterial lipopolysaccharide were dissolved in 100 μL of 25 mM EDTA, pH 7.5, with 1 mg/mL diastase (α-amylase and β-amylase) and 1 mg/mL lysozyme. The reaction was incubated at 50 °C for 30 minutes. After incubation, the reaction was filtered using an Microcon® - 10 Centrifugal Filters, 10 kDa NMWL (14,000 g, 30 minutes). To the fraction containing low molecular mass molecules (filtrate collected in the bottom tube), 50 μL of reaction buffer containing 5 mmol/L HD ligand was added, and saccharides were derivatized at 50°C for 20 minutes as described above.

### Acidic Digestion of Oligo- and Lipopolysaccharides

Saccharides and lipopolysaccharides were digested by formic acid and its isotopic variant (^13^C). One milligram of the sample was dissolved in 10 μL concentrated formic acid or formic acid-13C and incubated at 98 °C for 10 minutes in the thermocycler with a heated lid (104 °C). After incubation, 50 μL of 5% pyridine in water was added. The samples were filtered using Microcon® - 10 Centrifugal Filters, 10 kDa NMWL (14,000 g, 30 minutes), and the bottom fraction (50 μL) containing digested saccharides was used for derivatization using HD ligand and analysis as described above.

### Analysis of Lipopolysaccharides from Bacterial Cells

The bacteria were obtained from the Czech National Collection of Type Cultures (National Institute of Health, Prague, Czech Republic): *E. coli* ATCC25922, *E. coli* O55:B5 CNCTC5874, *E. coli* O111:B4 CNCTC5650, *Salmonella enterica* subsp. *enterica* serovar Enteritidis CNCTC5187, *S. enterica* subsp. *enterica* serovar Montevideo CNCTC6279, *S. enterica* subsp. *arizonae* CNCTC6478, *Shigella dysenteriae* CNCTC5204, Shigella boydii CNCTC6338, *Shigella flexneri* serovar 1a I:2,4 CNCTC6370, *Shigella flexneri* CNCTC6378 serovar 4a IV:3,4. *Legionella pneumophila* serogroups 1 (sequence type (ST) 1, ST23, ST62), and 10 (ST378) were obtained from our collection at the Department of Microbiology, Faculty of Medicine and University Hospital in Pilsen, Czech Republic. All members of the Enterobacterales order were cultivated on Mueller-Hinton agar at 35 °C overnight. Legionella pneumophila was cultivated on BCYE agar in an atmosphere with 5% CO2 for 24 hrs. The full loop of the bacteria was resuspended in 1 mL of 96% ethanol, centrifuged, and the pellet was allowed to dry at 98°C for 5 minutes. Twenty microliters of concentrated formic acid were added to the pellet and incubated at 98°C with shaking (1200 rpm) for 15 minutes. After incubation, 100 μL of 5% pyridine was added to adjust the pH to 3.0. The mixtures were then filtered using an Microcon® - 10 Centrifugal Filters, 10 kDa NMWL (14,000 g, 30 minutes), and the bottom filtrate (50 μL) was used for derivatization using HD ligand as described above. For microbes, further purification of derivatized saccharides is not required. Therefore, one microliter of the reaction mixture was directly applied to the MALDI target.

## RESULTS

### Synthesis of HD Ligand

Initially, we tested the derivatization of saccharides using rosaniline. Despite the very efficient binding of aldoses to the rosaniline, the disadvantage of the molecule was the presence of two efficient binding sites (NH_2_) and the requirement for a matrix for MALDI-TOF MS analysis. Therefore, we had tested several molecules containing an aldehyde residue to block one of those amines. Using vanillin, we obtained a molecule with an excellent ionization ability that does not require using any matrix in MALDI-TOF MS - an LDI-TOF (laser desorption/ionization time-of-flight) MS approach. It was, however, necessary to enhance the stability of the molecule by reductive amination of Schiff’s base (Supplementary Figure 1). Since a conjugated system is essential for efficient molecule ionization, we have tested an appropriate reducing agent (e.g., sodium cyanoborohydride, borane pyridine complex, 2-methylpyridine borane complex). Finally, we have chosen a 2-methylpyridine borane complex as the reaction can proceed in one step. It is also important to note that high concentrations of reducing agents, as recommended for conventional reactions, lead to the formation of leuco base in rosaniline. That molecule has very low ionization ability and requires a classical matrix-based MALDI-TOF MS setup.

The purity of the HD ligand synthesized in our laboratory was determined using LC-MS, and the structure was verified by solariX magnetic resonance mass spectrometry (Bruker Daltonics, Bremen, Germany) (Supplementary Figure 2). The quality of commercially synthesized HD ligand available as a 2,3-dichloro-5,6-dicyano-1,4-benzoquinone salt (DDQ) was determined by nuclear magnetic resonance (Supplementary Figure 3). Both variants showed equal results in the following experiments.

### Derivatization of Mono- and Disaccharides

The reaction was initially optimized using glucose and lactose. The best results were obtained in a low pH (3 - 4) and a buffer not containing chlorine ions. Therefore, we selected a pyridine buffer of pH 3.0 for further experiments. Using a MALDI-TOF mass spectrometer, the HD ligand’s ionization ability was excellent for detecting relevant signals of the conjugate with saccharides (Figure 1) without using any matrix (LDI-TOF MS). Ionization was performed in a positive ion mode with the analytes detected as [M+H]^+^ ions. All the saccharides tested provide a signal with a mass-to-charge ratio that can be calculated by the following formula (see Table 1):

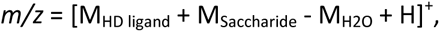

**Table 1.** LDI-TOF MS and LC-MS analysis of mono-, disaccharides, and 5-(hydroxymethyl)furfural showing *m/z* and ion mobility of derivatized molecules.

| Saccharide | $m/z$ | Ion<br>Mobility<br>[V.s/cm<sup>2</sup>] |
| --- | --- | --- |
| D-glucose | 586 | 1.242 |
| L-glucose | 586 | 1.242 |
| D-glucose-1,2-<sup>13</sup>C<sub>2</sub> | 588 | 1.242 |
| glucose-6-phosphate | 666 | 1.219 |
| L-rhamnose | 570 | 1.224 |
| L-fucose | 570 | 1.228 |
| D-xylose | 556 | 1.210 |
| lactose | 748 | 1.503 |
| maltose | 748 | 1.490 |
| 5-(hydroxymethyl)furfural | 532 | 1.099 |

Where the M_HD_ ligand is 423.195 (monoisotopic mass), M_H2O_ represents the loss of water (−18.015) during the formation of a Schiff base.

As demonstrated by glucose-6-phosphate, derivatization with HD ligand enables the detection of molecules with strongly negatively charged groups like phosphates in a positive ion mode as well (Figure 1). A purification step using C18 resin can be performed using the resin itself (as described in the methodology) or by ZipTips®, which enhances ionization ability in LDI-TOF MS analysis of mono- and oligosaccharides. This step removes unbound sugars that otherwise form a “sweet spot” and decreases the method’s sensitivity.

In LC-MS analysis, derivatized mono-, and di-saccharides have also been detected at the expected mass-to-charge ratio (Table 1, Figure 2). The ion mobility of the saccharides allows their further analysis and identification (e.g., discrimination between a disaccharide, lactose, and maltose – see Figure 2). Unfortunately, we could not distinguish between D- and L-glucose by our instrument.

**Figure 2.**
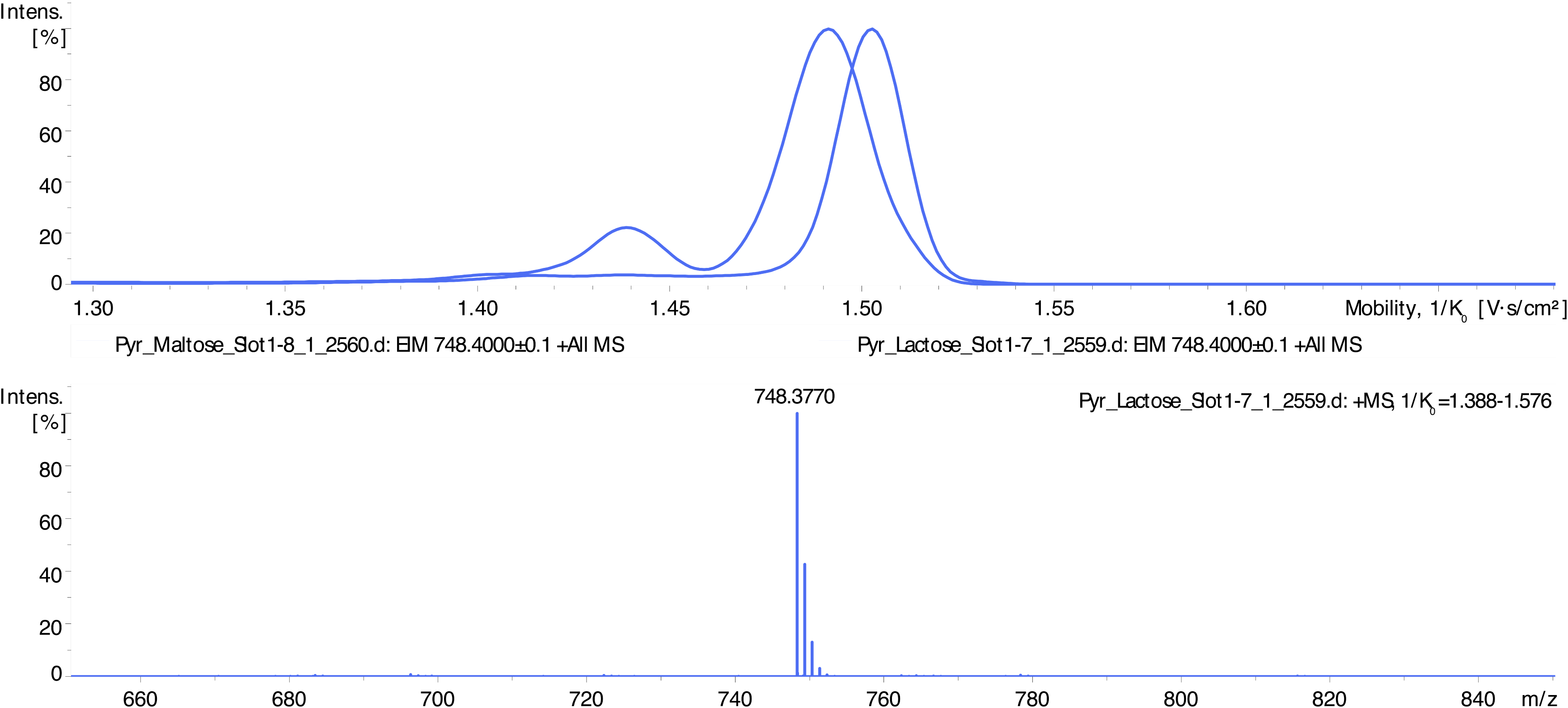
LC-MS spectra showing differentiation of disaccharides’ isoforms by ion mobility determination. Disaccharides maltose and lactose appear as signals at *m/z* 612 (HD ligand with the lost vanillin molecule) and *m/z* 748 (intact HD ligand). Maltose complex (*m/z* 748) shows an ion mobility of 1.490 V.s/cm^2^ and lactose 1.503 V.s/cm^2^.

The method’s sensitivity was tested using diluted glucose and lactose (0.01 mmol/L—100 mmol/L). For both methods (LDI-TOF MS and LC-MS), the sensitivity was determined to be 0.1 mmol/L.

### Acidic Digestion and Fischer-Speier Esterification

Using acidic digestion of oligosaccharides, we identified peaks that did not correspond with simple derivatized mono- and oligosaccharides. Using monosaccharides (_D_-glucose, _L_-fucose, _D_-xylose), formic acid, and isotopic formic acid-^13^C, the peaks corresponding to esters formed in hydroxyl groups of saccharides can be detected. In this reaction (Fischer-Speier esterification), the peaks are shifted by 28 g/mol, corresponding to formic acid-derived esters (Figure 3).

**Figure 3.**
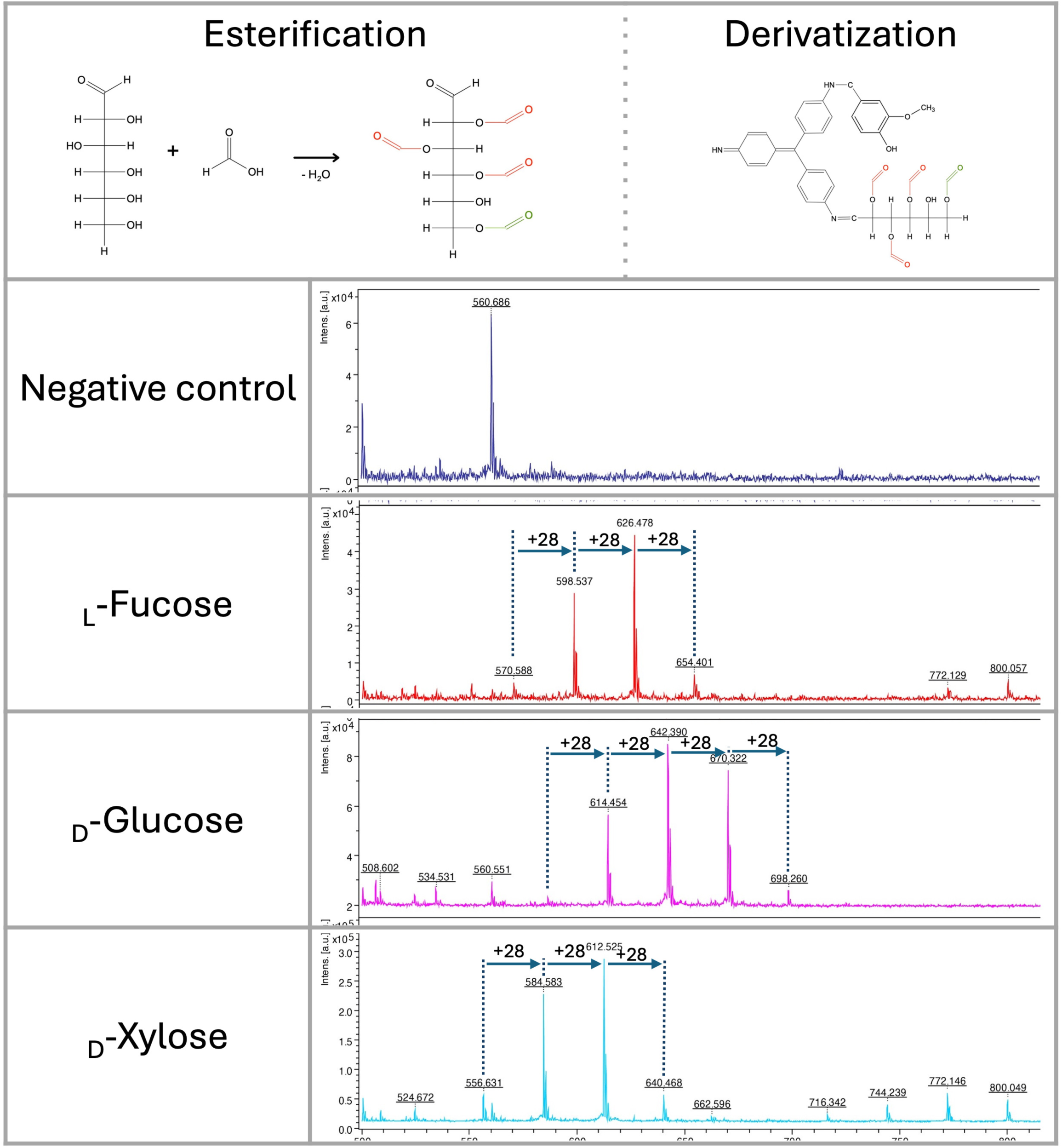
Fischer-Speier esterification of glucose using concentrated formic acid at 98 °C and its derivatization by HD ligand. Esterification of the hydroxyl group at position 6 (green color) was immediately detected. Esterification of other positions was identified in different ratios. LDI-TOF MS spectra of monosaccharides (negative control, _L_-fucose, _D_-glucose, _D_-xylose) esterified by concentrated formic acid (+*m/z* 28).

In glucose, the hydroxyl group at position 6 is almost completely esterified (*m/z* 614) with a very low signal of derivatized native glucose (*m/z* 586) (Figure 3). All except one hydroxyl group are also esterified with a different ratio, showing signals at *m/z* 642, 670, 698. Similar results were obtained in _D_-xylose and _L_-fucose, showing no signal with a non-esterified derivatized saccharide (*m/z* 556 and 570). As observed in glucose, variants of all esterified hydroxyl groups except one could be detected in LDI-TOF spectra (Figure 3). Based on those characteristics, we hypothesize that only one hydroxyl group of vicinal diols can be esterified efficiently. The esterification mechanism on D-glucose, D-xylose, L-fucose, and lactose was confirmed by precise molecular mass determination (<1 ppm) using solariX XR FT-ICR mass spectrometer (Supplementary Figure 4).

As hexoses in acidic conditions and high temperatures can be dehydrated to form 5-(hydroxymethyl)furfural, we also focused on identifying this molecule in the reaction. The 5-(hydroxymethyl)furfural can also be recognized as a signal at *m/z* 532 in acidic conditions. In glucose and fructose, the 5-(hydroxymethyl)furfural intermediate formed during saccharide dehydration [14] was identified in the spectra at *m/z* 550 (Figure 4, Figure 5).

**Figure 4.**
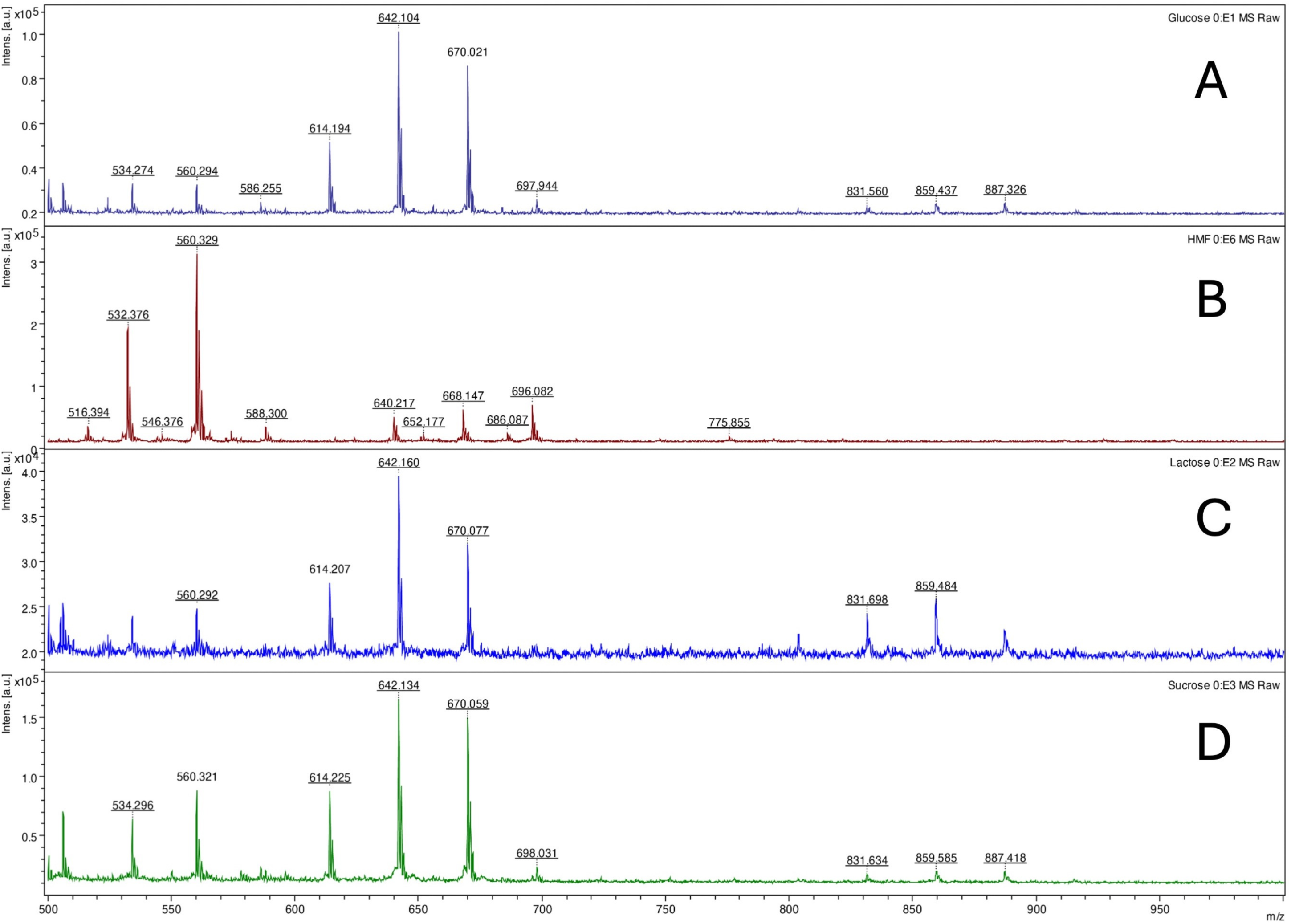
LDI-TOF MS spectra of glucose (A), 5-(hydroxymethyl)furfural (B), lactose (C), and sucrose (D) after digestion (lactose, sucrose) and esterification using concentrated formic acid at 98 °C for 10 minutes and derivatization using HD ligand.

**Figure 5.**
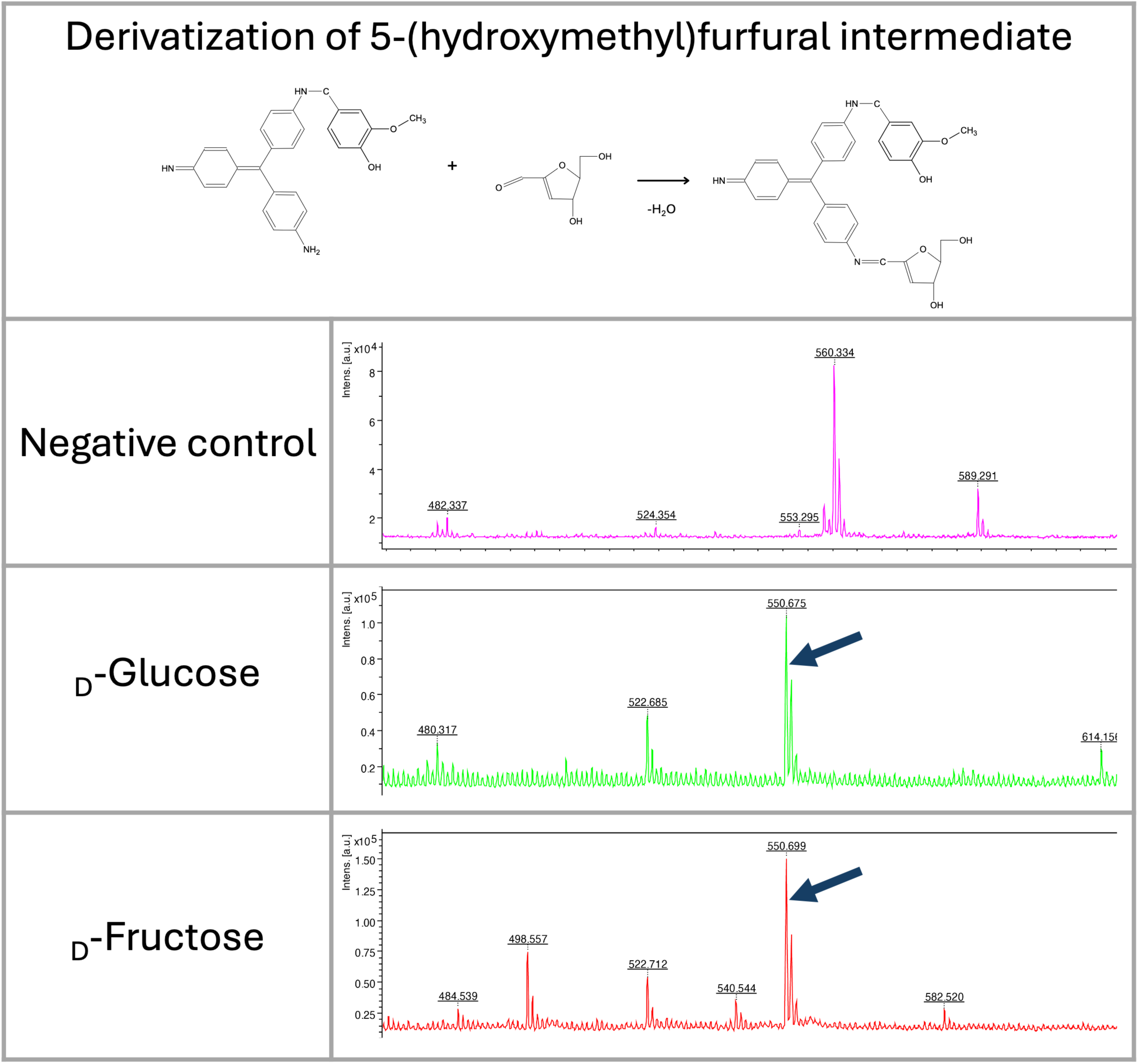
LDI-TOF MS spectra show the formation of 5-(hydroxymethyl)furfural intermediate from fructose and glucose after heating at 98 °C for 30 minutes (*m/z* 550). The molecule was derivatized by HD ligand and measured using LDI-TOF.

### Analysis of Bacterial Polysaccharides by Acidic Digestion

Initially, formic acid was tested for non-specific fingerprinting of polysaccharides. As demonstrated in Figure 4, concentrated hot formic acid (98 °C) can efficiently hydrolyze glycosidic bonds. The same results were obtained in raffinose and starch (data not shown). The method also allows the detection of common saccharides of the bacterial cell wall, a muramic acid, containing an amine and carboxyl group (*m/z* 657) (Figure 6).

**Figure 6.**
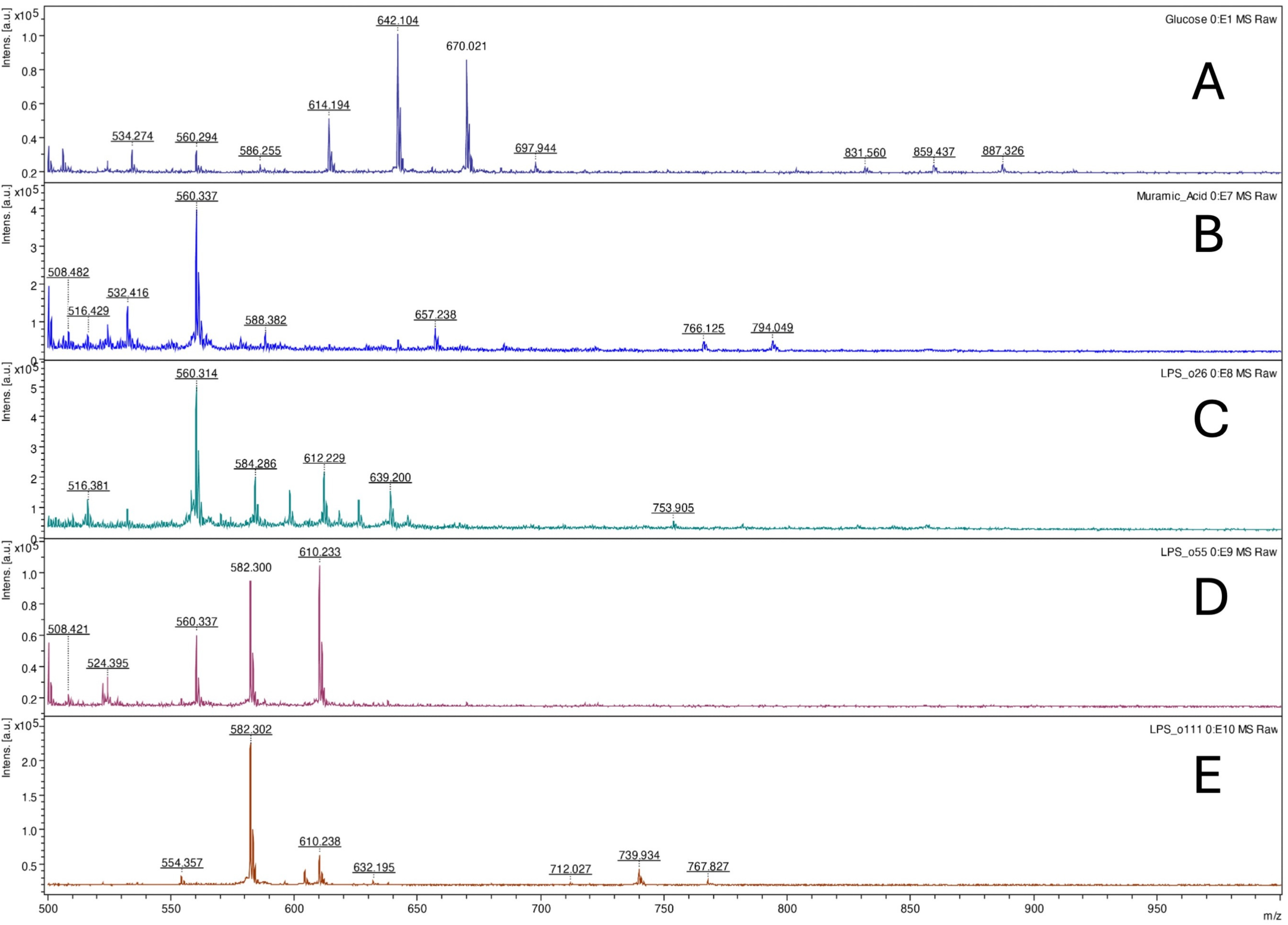
LDI-TOF MS spectra of glucose (A), muramic acid (B), *Escherichia coli* O26:B6 lipopolysaccharide (C), *E. coli* O55:B5 lipopolysaccharide (D), and *E. coli* O111:B4 lipopolysaccharide (E) after digestion and esterification using concentrated formic acid at 98 °C for 10 minutes and derivatization using HD ligand.

Based on those mono- and oligosaccharide results, the method was tested for fingerprinting bacterial lipopolysaccharides. Comparing three different bacterial polysaccharides of *Escherichia coli*, the results demonstrated different patterns. The same results were obtained for *Escherichia coli* O26:B6, E. coli O55:B5, E. coli O111:B4 purified lipopolysaccharides, and crude bacteria (Figure 7). Similarly, all tested bacteria, including *Salmonella* spp., *Shigella* spp., and *Legionella pneumophila*, provided specific patterns showing that the method can be used for bacterial typing.

**Figure 7.**
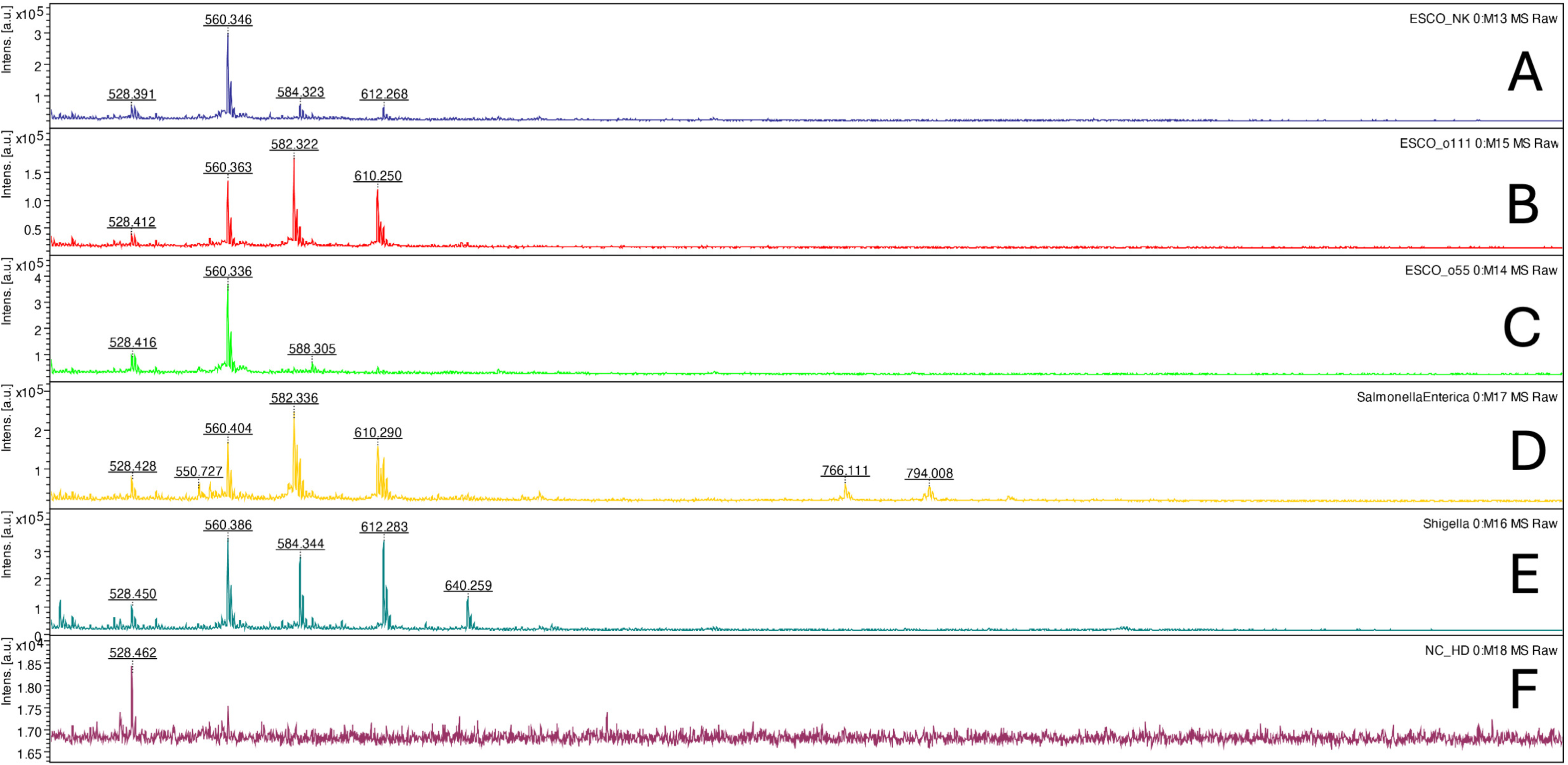
LDI-TOF MS spectra of *Escherichia coli* ATCC25922 (A), *E. coli* O111:B4 CNCTC5650 (B), *E. coli* O55:B5 CNCTC5874 (C), *Salmonella enterica* subsp. *enterica* serovar Enteritidis CNCTC5187 (D), *Shigella dysenteriae* CNCTC5204 (E), and negative control (F) after digestion and esterification using concentrated formic acid at 98 °C for 10 minutes and derivatization using HD ligand.

### Analysis of Polysaccharides by Enzymatic Digestion

To analyze polysaccharides and lipopolysaccharides, α-amylase and β-amylase were tested using starch as a positive control. In starch, mono—(*m/z* 586), di—(*m/z* 748), and trisaccharide (*m/z* 910) can be observed using both methods (LDI-TOF MS and LC-MS). We also detected distinct lipopolysaccharide profiles (*E. coli* O26:B6, *E. coli* O55:B5, *E. coli* O111:B4) similar to acidic digestion.

## DISCUSSION

We describe here a novel method for derivatizing mono- and oligosaccharides that can be used for the identification and analysis of those molecules, not only restricted to microbial origin. Initially, we focused on developing a technique to identify bacterial cell-wall polysaccharides (i.e., lipopolysaccharides). We tested several derivatizing agents, as saccharides cannot be easily ionized compared to peptides/proteins by MALDI-TOF mass spectrometry. Inspired by a detection of lactose fermentation using a basic fuchsin in diagnostic bacteriology (e.g., Endo agar), we tested that molecule to form a Shiff base between the aldehyde group of reducing sugars and the amine of the fuchsin. This complex could be derivatized by MALDI-TOF MS using a standard matrix (e.g., 2,5-dihydroxybenzoic acid). The disadvantage of this process is the presence of two efficient amine residues in the molecule, responsible for the polyvalent binding of tested saccharides. However, the ability to modify saccharides with fuchsin and subsequent ionization showed excellent results. Therefore, we decided to focus our research further on modifying this molecule.

After modifying the fuchsin by adding vanillin to one of the amines, a designated HD ligand, allowed the ionization of the complex with saccharides without using the matrix (LDI-TOF MS). The complex can also be analyzed using LC-MS with a separation on the C18 reverse phase column. As demonstrated in the results (Figure 1), some isomeric saccharides can also be distinguished by their ion mobility using trapped ion mobility technology.

The HD ligand allows the derivatization and analysis of saccharides with different substituents, including phosphates, in a positive ion mode. This makes the method universal to detect microbial oligo- and polysaccharides of different origins. However, it is necessary to use optimal reaction conditions for derivatization, including a pH between 3 and 5. We also found that a high concentration of chlorine ions inhibits the formation of a Schiff base (data not shown). Among the buffers tested, pyridine in a concentration between 1 and 10% with a pH adjusted by formic or acetic acid provided the highest binding efficiency.

Although we expected that the fuchsin-based system would also allow the detection of ketones (ketoses), we could not find conditions that would enable efficient, stable binding of the HD ligand. This may be because the HD ligand requires specific binding conditions or is unstable upon ionization during mass spectrometry. On the other hand, however, ketoses are not common structures in bacterial cells. For analysis, they can be modified to furfurals in acidic conditions that are analyzable by our system as well (see Figure 5).

When different options for the hydrolysis of the glycosidic bond of polysaccharides were tested, we found that using formic acid, spectra containing many ions of different m/z with regular repetitions (+28) were detected. By detailed analysis, including the reaction in 13C formic acid, we identified the signals as Fischer-Speier esters formed in the hydroxyl groups of saccharide molecules. This behavior can be further used to identify and analyze saccharides more accurately. For future experiments, the relative intensity of signals representing esterified hydroxyl groups can be further verified to determine their position in the molecule. We also tested acetic acid for esterification. However, its efficiency was very low compared with formic acid. In glucose, an acetic acid-derived ester was formed at position six only (data not shown). Those findings were crucial for further microbial polysaccharide experiments to understand the reaction.

An essential step in the analysis of microbial polysaccharides is to digest the molecule specifically, which usually possesses a very high molecular weight. Similarly to derivatization methods, we tested many possibilities, including specific lipopolysaccharide isolation, e.g., Bligh-Dyer solution and its modifications (data not shown). Interestingly, common amylases (α and β) can efficiently digest bacterial polysaccharides to mono-, di-, and trisaccharides. Peptidoglycan-specific enzymes (e.g., lysozyme) can also be used for specific analysis. A similar approach can be optimized to analyze other microbial polysaccharides (e.g., galactomannan and glucan in molds and yeasts) that are clinically relevant for rapidly diagnosing invasive fungal infection or detecting resistance to antifungal drugs.

Finally, we selected a straightforward method that does not require any specific extraction of cell-wall polysaccharides: incubating microbes in concentrated formic acid at 98 °C. This procedure blocks reactive amines in the crude microbes, and the polysaccharides can be simultaneously digested into mono and oligosaccharides in a one-step process without previous specific extraction of the cell walls, which is usually laborious. That approach is a typical example of applying Ockham’s razor and can be easily used in routine diagnostic laboratories.

In all applications, however, it is crucial to filter the reaction mixture before the derivatization of saccharides to remove undigested polysaccharides (> 3 – 10 kDa). Without this step, the spectra show an insufficient noise-to-signal ratio.

## CONCLUSIONS

The method described here can be used to analyze microbial polysaccharides not only for epidemiological typing and polysaccharide vaccine development but also to open the possibility of detecting those structures directly in clinical specimens. Derivatization of saccharides and related molecules (e.g., furfurals) containing an aldehyde group using the HD ligand is challenging for further analysis of those important biological structures by mass spectrometry (LDI-TOF MS, LC-MS).

## SUPPLEMENTARY MATERIAL

- **Supplementary Figure 1** - Principle of low-scale preparation of Vanillyl-Rosaniline (HD) ligand using 2-methylpyridine borane complex as a reduction agent and purification by HPLC system.
- **Supplementary Figure 2** - Confirmation of Vanillyl-Rosaniline (HD) ligand structure using 15T solariX XR FT-ICR mass spectrometer (Bruker Daltonics).
- **Supplementary Figure 3** - Quality control protocol (NMR spectra) of commercially prepared Vanillyl- Rosaniline (HD) ligand.
- **Supplementary Figure 4** - Confirmation of esterified _D_-glucose, _D_-xylose, _L_-fucose, and lactose using 15T solariX XR FT-ICR mass spectrometer (Bruker Daltonics).

## AUTHOR CONTRIBUTIONS

L.D. was responsible for laboratory work, interpretation of the results, designing novel derivatization agents, and writing the manuscript. V.P. was responsible for laboratory work, the result interpretation, and the manuscript’s writing. P.N. was responsible for confirming HD ligand structure and analysis of esterified products. J.H. was responsible for conceptualization, methodology, designing new derivatization agents, data interpretation, and writing the manuscript.

## ACKNOWLEDGMENT

We thank Dana Kralova and Andrea Pospechova for their excellent technical assistance. The study was supported by the Czech Health Research Council grant Nr. NW24-09-00464, the Charles University Grant Agency (GA UK) project Nr. 550225, and the National Institute of Virology and Bacteriology (Programme EXCELES, ID Project No. LX22NPO5103)—funded by the European Union—Next Generation EU. The method has been patented, including the HD reagent molecule and its variants (Czech National Patent PV 2024-48, PCT/CZ2025/050014).

